# Enlarged Perivascular Spaces are Associated with White Matter Injury, Brain Atrophy, Cognitive Decline and Markers of Inflammation in an Autosomal Dominant Vascular Neurodegenerative Disease (CADASIL)

**DOI:** 10.1101/2023.08.17.553732

**Authors:** Nikolaos Karvelas, Bradley Oh, Earnest Wang, Yann Cobigo, Torie Tsuei, Stephen Fitzsimons, Alexander Ehrenberg, Michael Geschwind, Daniel Schwartz, Joel Kramer, Adam R. Ferguson, Bruce L. Miller, Lisa Silbert, Howard Rosen, Fanny M. Elahi

## Abstract

**Background and Objectives:** Enlarged perivascular spaces (ePVS) have been previously reported in Cerebral Autosomal Dominant Arteriopathy with Subcortical Infarcts and Leucoencephalopathy (CADASIL), but their significance and pathophysiology remains unclear. We investigated associations of ePVS with classical imaging measures, cognitive measures and plasma proteins to better understand what ePVS represents in CADASIL and whether radiographic measures of ePVS would be of value in future therapeutic discovery studies for CADASIL.

**Methods:** 24 individuals with CADASIL and 24 age and sex matched controls were included. Disease status was determined based on presence of *NOTCH3* mutation. Brain imaging measures of white matter hyperintensity (WMH), brain parenchymal fraction (BPF), ePVS volumes, clinical, and cognitive measures, as well as plasma proteomics were used in models. Global ePVS volumes were calculated via a novel, semi-automated pipeline and levels of 7363 proteins were quantified in plasma using the SomaScan assay. The relationship of ePVS with global burden of WMH, brain atrophy, functional status, neurocognitive measures, and plasma proteins were modelled with linear regression models.

**Results:** CADASIL and control groups did not exhibit differences in mean ePVS volumes. However, increased ePVS volumes in CADASIL were associated with increased WMH volume (β=0.57, p=0.05), Clinical Dementia Rating (CDR) Sum-of-Boxes score (β=0.49, p=0.04), and decreased brain parenchymal fraction (BPF) (β=-0.03, p=0.10). In interaction term models, the interaction term between CADASIL disease status and ePVS volume was associated with increased WMH volume (β=0.57, p=0.02), Clinical Dementia Rating (CDR) Sum-of-Boxes score (β=0.52, p=0.02), decreased BPF (β=-0.03, p=0.07) and Mini Mental State Examination (MMSE) score (β=-1.49, p=0.03). Proteins positively associated with ePVS volumes were found to be related to leukocyte migration and inflammation, while negatively associated proteins were related to lipid metabolism. Two central hub proteins were identified in protein networks associated with ePVS volumes: CXCL8/IL-8, and CCL2/MCP-1. The levels of CXCL8/IL8 were also associated with increased WMH volume (β=2.44, p < 0.01), and levels of CCL2/MCP-1 were further associated with decreased BPF (β=-0.0007, p < 0.01), MMSE score (β=-0.02, p < 0.01), and increased Trail Making Test B (TRAILB) completion time (β=0.76, p < 0.01). No protein was associated with all 3 studied imaging measures of pathology (BPF,ePVS,WMH).

**Discussion:** Based on associations uncovered between ePVS volumes and cognitive functions, imaging and plasma proteins, we conclude that ePVS volumes capture pathologies contributing to chronic brain dysfunction and degeneration in CADASIL, with relevance to future clinical trials for novel therapeutic discoveries to prevent decline and injury in individuals carrying *NOTCH3* mutations.

## Introduction

Considering the high prevalence of cerebral small vessel disease (cSVD), most commonly as incidental imaging findings^1, 2^, studies of vascular contributions to cognitive impairment and dementia (VCID) represent a critical area of research. A toolbox of biomarkers for accurate quantification of cSVD *in vivo* is needed for design of impactful clinical trials and therapeutic advances^2^. Cerebral Autosomal Dominant Arteriopathy with Subcortical Infarcts and Leukoencephalopathy (CADASIL), a monogenic form of VCID^3^, provides a unique opportunity for modeling molecular and structural pathologies for identification of biological signatures and therapeutic targets across the spectrum of VCID disease severity.

CADASIL, primarily caused by mutations in the epidermal growth factor–like repeats (EGFr) of the *NOTCH3* gene, is the most common monogenic form of vascular neurodegenerative disease, with migraines, strokes, neuropsychiatric symptoms, and cognitive impairment emerging by midlife^3^. In early stages, radiographic markers can be abnormal even in the absence of significant clinical symptoms^4^. White matter hyperintensities (WMH) on Fluid Attenuated Inversion Recovery (FLAIR) magnetic resonance imaging (MRI), a classic imaging biomarker of CADASIL, is detectable years before symptom onset^4-6^, however WMH do not consistently track with symptomatic progression of cognitive dysfunction^7-9^. Similarly, cerebral microbleeds (CMBs), another characteristic radiographic finding, albeit less prevalent than WMH, remain on brain imaging life-long and may therefore not track the dynamic progression in disease^7, 10^. Amongst radiographic biomarkers of cSVD in CADASIL, lacunes have demonstrated the most consistent association with progressive cognitive decline ^9, 11, 12^, although they occur in later disease stages, past the earliest desirable windows for therapeutic interventions^3^.

Perivascular spaces (PVS) are cerebrospinal fluid (CSF) filled spaces that surround small arterioles and venules of the brain. PVS represent a major clearance route for the brain, as well as a microenvironment where immune cells and glia interact with blood vessels^13^. Not all PVS are visible on brain imaging. The radiographically visible and quantifiable PVS are referred to as enlarged perivascular spaces (ePVS) in *in vivo* human studies^14^. MRI visible ePVS follow the typical course of vessels and appear isointense to CSF^14^. It is thought that perivascular space enlargement can reflect structural and functional changes in cerebral microvessels and can represent the accumulation of stagnant CSF, perivascular immune cells, metabolites, and proteins, including toxic substrates such as amyloid beta, in the glymphatic system along vessels^15, 16^. Pathophysiological mechanisms that have are thought to influence ePVS development are increased vascular tortuosity related to dysregulated angiogenesis, changes to elasticity, blood-brain barrier dysfunction, immune activation, and changes to the extracellular matrix, as well as multi-factorial clearance defects^17^.

In CADASIL, ePVS remain relatively understudied and may represent a relevant biomarker that predicts cognitive decline, while capturing therapeutically interesting pathologies for future interventional trials. In this study we use a rigorous quantitative method for measuring whole brain ePVS volumes to examine differences between individuals carrying *NOTCH3* mutations and age and sex matched controls. We then test associations with clinically-meaningful imaging and cognitive outcomes. Finally, we provide insights regarding possible implicated molecular mechanisms through associations with plasma proteomics.

## Methods

### Study Participants

Consecutive enrollment of CADASIL patients (n=24) was undertaken from February 25, 2019 to August 2, 2021 into our prospective longitudinal study of CADASIL, VascBrain. The study was based at the UCSF Memory and Aging Center and is currently at the Icahn School of Medicine at Mount Sinai. CADASIL was confirmed based on sequencing of *NOTCH3*. Age and sex matched controls (n=24) were sampled from ongoing longitudinal studies at the Memory and Aging Center at UCSF (Chronic Inflammation study, MarkVCID, Larry J. Hillblom foundation study, ARTFL-LEFFTDS Longitudinal Frontotemporal Lobar Degeneration Study and the MAC Alzheimer’s Disease Research Center study). All controls were selected based on normal functional status, absence neurological disease assessed by history and physical exam, or findings concerning for CADASIL. Exclusion criteria of parent studies applied to this study, including active or uncontrolled psychiatric disease such as psychosis, brain tumor, or history of brain surgery. No exclusion criteria based on imaging or cognitive measures were defined, so that controls better represented the general population.

### Clinical and Cognitive Evaluation

A brief health history and physical examination were noted for all participants. Participants were also given a standard battery of clinical and neuropsychological tests. Clinical Dementia Rating (CDR) and CDR sum of boxes (CDR-SB) were completed on all participants via study partner interviews. For our analyses, we included CDR as a surrogate of functional impairment and measures of general cognition (Mini Mental State Examination - MMSE), executive function (time to complete Trail Making Test B - TRAILB), and episodic visual memory (modified Rey-Osterrieth Complex Figure delayed recall at 10 minutes - Rey10m).

### Neuroimaging

Participants had MRI acquired on a Siemens Tim Trio 3 Tesla scanner using the local protocol, which acquired T1-weighted imaging using an MP-RAGE sequence with the following parameters: 160×240×256 matrix; 160 slices; voxel size = 1×1×1 mm^3^; flip angle 9°, TE 2.98 ms, TR 2300 ms. The T2-weighted used the following parameters: 176×256×256 matrix; 176 slices; voxel size = 1×1×1 mm^3^; TE 408 ms, TR 3200 ms. The FLAIR sequence was acquired with slice thickness = 1.00mm; slices per slab = 176; in-plane resolution = 1.0×1.0mm; matrix = 256×256; TR = 5000ms; TE = 397ms; TI = 1800ms; flip angle = 120°.

Before preprocessing of the images, all T1-weighted images were visually inspected for quality. Images with excessive motion or other artifacts were excluded. T1-weighted images underwent bias field correction using the N3 algorithm, and segmentation was performed using the SPM12 (Wellcome Trust Center for Neuroimaging, London, UK) unified segmentation^18^. A group template was generated from subject gray and white matter tissues and cerebrospinal fluid using the Large Deformation Diffeomorphic Metric Mapping framework^19^. Every step of the transformation was carefully inspected from the native space to the group template.

Brain Parenchymal Fraction (BPF) was calculated as the quotient of parenchymal tissue volume by total intracranial volume (TIV) and represents a measure of brain atrophy. Global cerebral volume of WMH was measured on T2/FLAIR sequences and was rated by a board-certified neurologist (FME) in addition to being reviewed by a neuroradiologist to rule out other significant abnormalities.

An unsupervised segmentation algorithm was used to quantify ePVS burden within white matter (WM) tissue. Neuroimaging standards established by Wardlaw et al.^14^ were used in the identification and analysis of ePVS. Total ePVS burden was quantified using a 90% binarized white matter mask created in the T1-weighted native space, where automatic fluid-filled areas with contrast resembling cerebrospinal fluid (CSF) were identified using FSL^20^. Corresponding T2-weighted images were registered into T1-weighted space. We enhanced the ePVS presence in the WM using the method developed in Boespflug et al.^21^. The proposed linear model used (1) T1- and T2-weighted images normalized by their mean values defined within the WM mask, which served as the response variable; and (2) T1 and T2 mean values calculated in the gray matter (GM) and CSF, which served as nuisance and predictor variables. Correlation coefficients were calculated from CSF fluid-filled regions identified in the WM mask, such that areas of high positive value were selectively filtered as true representations of ePVS. We used an additional method, implemented in ITK, to selectively identify tortuous, vessel-like geometries from segmented, selectively-filtered ePVS^22^, by approximating the eigenvalues of the Hessian matrix at each voxel. The method factored in the modulation of eigenvalues along the principal direction of each identified ePVS to selectively filter regions of true vessel-like enlargement. An optimized Sato filter was then applied to differentiate tortuous vessel geometries of ePVS from ovoid lacunar infarcts. A density-based clustering algorithm iterated on the segmented mask to independently label unique ePVS, providing an estimate of total burden.

Visual assessment of clustered segmentation output was performed by three imaging experts (BO, EW, TT) to manually exclude false positives, including anatomical boundaries resembling tortuous geometries (e.g. sulci, cingulate, ventricles, and tissue boundaries between deep gray matter regions) and other manifestations of small vessel disease (e.g. lacunar infarcts, ischemic stroke infarcts, and severe white matter disease), as well as false negatives, including ePVS in granular subcortical areas such as the basal ganglia.

### Plasma Collection and Proteomic Analysis

Blood samples were collected in EDTA tubes from all CADASIL patients. Samples were then centrifuged at 1000g for 10 minutes, then at 2500g for 10 minutes to obtain platelet-poor plasma and stored at -80°C. Afterwards they were sent to be analyzed with the SOMAscan 7k assay (SomaLogic, Inc., Boulder, CO) according to standardized protocols described elsewhere^23^. Briefly, the SOMAscan assay kit employs 7596 highly selective single-stranded modified Slow Off-rate Modified DNA Aptamers (SOMAmer) for protein identification and quantification.

Through the use of a custom DNA microarray (Agilent), data is reported as relative fluorescence units (RFU). SomaLogic performed quality control, calibration and normalization of the data, and analysis of all samples was deemed appropriate. Non-human SOMAmers were removed from the dataset, leaving 7363 relevant proteins for further analysis.

For the protein analyses, we performed outlier analyses with the interquantile range (IQR) method and removed 2 individuals from the CADASIL group with ePVS volumes over 1.5 times the IQR >75th percentile (Q3). We inputted proteins of interest in the STRING database version 12.0 for protein-protein functional and physical interaction analysis, the results of which were displayed as a functional network. Interactions were considered with a medium confidence score of 0.4 or higher. Biological process Gene Ontology (GO) analysis was performed through the clusterProfiler R package^24^, which implements the Bioconductor *Homo sapiens* annotation package org.Hs.eg.db.

### Statistical Analysis

Mean demographics, imaging, and cognitive test measures were compared between CADASIL and control cohorts with Student’s t-test for continuous variables, Mann-Whitney U-test for nominal variables and Chi-square tests for categorical variables. Linear regression models were used to test for associations. For the regression models, WMH and ePVS volume were standardized in the whole sample to have a mean of 0 and a SD of 1. These models were then adjusted for age, sex and education. A different set of linear regression models was performed with the interaction between ePVS and the presence or absence of CADASIL (disease status) as the main predictor. Interaction model lines were fitted in interaction plots. Linear regression models were used to examine associations between the measures of interest and proteomic markers. Variance inflation factors (VIF) for the covariates of multivariate models were inspected to test for multicollinearity. All analyses were performed using R, version 4.2.1 (R Foundation for Statistical Computing, 2022). All statistical tests were unpaired. A 2-sided p-value ≤ 0.05 was considered statistically significant, and a p-value < 0.10 but greater than 0.05 marginally significant.

### Standard Protocol Approvals, Registrations, and Patient Consents

Study protocols were approved by the UCSF Human Research Protection Program and Institutional Review Board. Research was performed in accordance with the Code of Ethics of the World Medical Association Written informed consent was obtained from all patients before data collection.

### Data Availability

All de-identified data employed in this study are available upon reasonable request from any qualified investigator for replication of procedures and results.

## Results

### Sample demographics and clinical characteristics

Participant characteristics are summarized in Table 1. The control group (n = 24) was selected to be age and gender-matched to the CADASIL patient group (n = 24). Mean (SD) age was 50.69 (12.45) years for the CADASIL patients and 52.8 (11.26) years for the control participants. 16 (67%) participants in each group were female. The control group had significantly more years of educational attainment (17.09, [2.13]) than the CADASIL group (14.7, [4.17]) (p = 0.02).

**Table 1.** Summary of demographic, clinical and imaging characteristics. Values are reported as mean ± standard deviation. Two-tailed Student’s t-tests or the non-parametric Wilcoxon rank sum were used for statistical comparison of continuous and nominal measures and Chi-squared tests were performed to compare group characteristics. *p-value < 0.05.

|  | CADASIL<br>(n = 24) | Control<br>(n = 24) | p-value |
| --- | --- | --- | --- |
| **Demographics** |  |  |  |
| Age, [range] | 50.69 ± 12.45<br>[23-75] | 52.82 ± 11.26<br>[28-75] | 0.54 |
| Sex, n (%) female | 16 (67%) | 16 (67%) | 1.0 |
| Education in years | 14.70 ± 4.17 | 17.09 ± 2.13 | **0.02\*** |
| **Cognitive Measures** |  |  |  |
| Clinical Dementia Rating (CDR)<br>Global Score | 0.17 ± 0.24 | 0.02 ± 0.10 | **0.01\*** |
| CDR Sum of Boxes | 0.54 ± 0.83 | 0.15 ± 0.50 | **0.01\*** |
| Mini Mental State Examination<br>(MMSE) Total Score | 27.72 ± 2.49 | 29.52 ± 0.59 | **0.02\*** |
| Trail B Test Time (TRAILB) (sec) | 84.67 ± 65.72 | 47.88 ± 14.69 | **0.02\*** |
| Rey-Osterrieth Complex Figure<br>Test recall at 10 minutes (Rey10m) | 12.24 ± 2.19 | 13.36 ± 2.08 | 0.09 |
| **Imaging Characteristics** |  |  |  |
| Brain Parenchymal Fraction (BPF) | 0.77 ± 0.06 | 0.78 ± 0.03 | 0.54 |
| Total White Matter Hyperintensity<br>(WMH) volume (mm<sup>3</sup>) | 12723.27 ±<br>8334.02 | 1306.34 ± 721 | **0.0000007\*** |
| Enlarged Perivascular Spaces<br>(ePVS) volume (mm<sup>3</sup>) | 547.75 ±<br>441.38 | 487.88 ±<br>747.55 | 0.73 |

Concerning cognitive features, CADASIL patients had higher scores in the CDR Global scale (0.17 [0.24] vs 0.02 [0.10]; p = 0.01) and the CDR Sum of Boxes score (0.54 [0.83] vs 0.15 [0.50]; p = 0.01), lower MMSE scores (27.72 [2.49] vs 29.52 [0.59]; p = 0.02), and took more time to complete the Trail B test (84.67 [65.7] vs 47.88 [14.7] seconds; p = 0.02). As expected, there was no significant difference for the delayed recall score of the modeified Rey - Osterrieth Complex Figure test (p=0.09), which was used as a surrogate of visual episodic memory.

Comparisons between imaging measures revealed no significant group differences with regards to brain parenchymal fraction (p=0.54) or ePVS volumes (p=0.73). Total WMH volume was significantly increased in the CADASIL group, as expected (12723 [8334] vs 1306 [721] mm^3^; p < 0.001).

### Associations of ePVS with Imaging and cognitive measures

Results from all models are presented in Table 2. In the unadjusted models, increase in standardized ePVS volume was associated with increase in standardized WMH volume (β = 0.57; 95% CI = 0.01 to 1.14; p = 0.05) and in the CDRBox score (β = 0.49; 95% CI = 0.03 to 0.95; p = 0.04) for CADASIL but not controls. We also found a marginally significant decrease in BPF (β = -0.03; 95% CI = -0.06 to 0.005; p = 0.10) with increased ePVS volume. In the models adjusting for age, sex, and education, a similar trend was observed for CDRBox (β = 0.41; 95% CI = 0.04 to 0.77; p = 0.03) but not for the other measures. Expectedly, no significant associations were found in unadjusted and adjusted models for the control group between standard ePVS volumes and any of the outcomes. Surprisingly, in the adjusted model, TRAILB completion time seemed to decrease with increased ePVS volume, but the effect was marginally significant (β = -19.72; 95% CI = -40.11 to 0.68; p = 0.06).

**Table 2.**
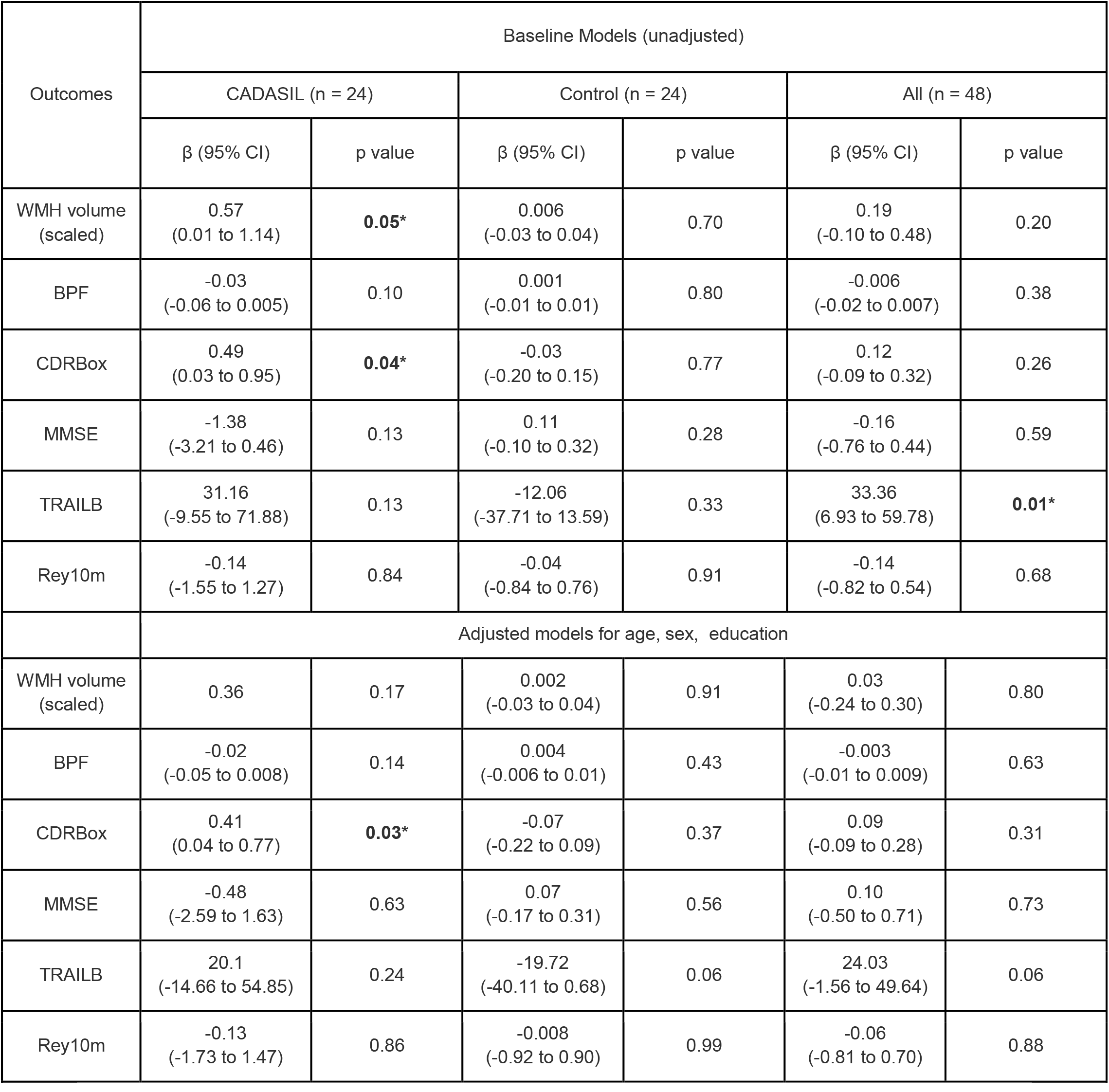
Baseline and adjusted linear regression model associations with standardized ePVS volume as main predictor and imaging and cognitive measures as outcomes. Abbreviations: BPF = Brain Parenchymal Fraction, CADASIL = Cerebral Autosomal Dominant Arteriopathy with Subcortical Infarcts and Leucoencephalopathy, CDRBox = Clinical Dementia Rating Sum-of-Boxes, MMSE = Mini Mental State Examination, Rey10m = Rey-Osterrieth Complex Figure Test delayed recall at 10 minutes, TRAILB = Trail B Test completion time, WMH = White Matter Hyperintensity. *p-value < 0.05.

| Outcomes | Baseline Models (unadjusted) |  |  |  |  |  |
| --- | --- | --- | --- | --- | --- | --- |
|  | CADASIL (n = 24) |  | Control (n = 24) |  | All (n = 48) |  |
| | $\beta$ (95% CI) | p value | $\beta$ (95% CI) | p value | $\beta$ (95% CI) | p value |
| WMH volume (scaled) | 0.57<br>(0.01 to 1.14) | **0.05\*** | 0.006<br>(-0.03 to 0.04) | 0.70 | 0.19<br>(-0.10 to 0.48) | 0.20 |
| BPF | -0.03<br>(-0.06 to 0.005) | 0.10 | 0.001<br>(-0.01 to 0.01) | 0.80 | -0.006<br>(-0.02 to 0.007) | 0.38 |
| CDRBox | 0.49<br>(0.03 to 0.95) | **0.04\*** | -0.03<br>(-0.20 to 0.15) | 0.77 | 0.12<br>(-0.09 to 0.32) | 0.26 |
| MMSE | -1.38<br>(-3.21 to 0.46) | 0.13 | 0.11<br>(-0.10 to 0.32) | 0.28 | -0.16<br>(-0.76 to 0.44) | 0.59 |
| TRAILB | 31.16<br>(-9.55 to 71.88) | 0.13 | -12.06<br>(-37.71 to 13.59) | 0.33 | 33.36<br>(6.93 to 59.78) | **0.01\*** |
| Rey10m | -0.14<br>(-1.55 to 1.27) | 0.84 | -0.04<br>(-0.84 to 0.76) | 0.91 | -0.14<br>(-0.82 to 0.54) | 0.68 |
|  | Adjusted models for age, sex, education |  |  |  |  |  |
| WMH volume (scaled) | 0.36 | 0.17 | 0.002<br>(-0.03 to 0.04) | 0.91 | 0.03<br>(-0.24 to 0.30) | 0.80 |
| BPF | -0.02<br>(-0.05 to 0.008) | 0.14 | 0.004<br>(-0.006 to 0.01) | 0.43 | -0.003<br>(-0.01 to 0.009) | 0.63 |
| CDRBox | 0.41<br>(0.04 to 0.77) | **0.03\*** | -0.07<br>(-0.22 to 0.09) | 0.37 | 0.09<br>(-0.09 to 0.28) | 0.31 |
| MMSE | -0.48<br>(-2.59 to 1.63) | 0.63 | 0.07<br>(-0.17 to 0.31) | 0.56 | 0.10<br>(-0.50 to 0.71) | 0.73 |
| TRAILB | 20.1<br>(-14.66 to 54.85) | 0.24 | -19.72<br>(-40.11 to 0.68) | 0.06 | 24.03<br>(-1.56 to 49.64) | 0.06 |
| Rey10m | -0.13<br>(-1.73 to 1.47) | 0.86 | -0.008<br>(-0.92 to 0.90) | 0.99 | -0.06<br>(-0.81 to 0.70) | 0.88 |

When combining both CADASIL and control participants into one group, an increase in ePVS volume was associated with slowed processing speed in the unadjusted model (β = 33.36; 95% CI = 6.93 to 59.78; p = 0.01). Furthermore, the association was marginally significant in the fully adjusted model (β = 24.03; 95% CI = -1.56 to 49.64; p = 0.06). No other associations emerged. No multicollinearity was detected for any of the covariates in the multivariate models (VIF < 5).

### Associations between ePVS - CADASIL interaction with imaging and cognitive measures

Since we observed associations mainly in the CADASIL group, we proceeded to examining interaction models run in both groups combined (n = 48). The values derived for the interaction term between disease status and ePVS volume (CADASIL status x ePVS volume) are presented in Table 3. Some of the interaction models are visualized as interaction plots of partial residuals in Figure 1. WMH volume was positively associated in both baseline (β = 0.57; 95% CI = 0.12 to 1.02; p = 0.015), and adjusted models (β = 0.45; 95% CI = 0.03 to 0.87; p = 0.03) (Figure 1a). BPF had a marginally significant negative association with the interaction term in the unadjusted (β = -0.03; 95% CI = -0.06 to -0.002; p = 0.07). (Figure 1b), and in the adjusted model (β = -0.02; 95% CI = -0.05 to 0.0007; p = 0.06) Regarding cognitive measures, we found CDRBox and MMSE scores to be significantly associated with the interaction term. Specifically, CDRBox scores increased as interaction term values increased in the baseline (β = 0.52; 95% CI = 0.08 to 0.95; p = 0.02),) and fully adjusted models (β = 0.39; 95% CI = 0.01 to 0.77; p = 0.04) (Figure 1c). MMSE scores declined as interaction term values increased in the baseline model (β = -1.49; 95% CI = -2.81 to -0.16; p = 0.03) (Figure 1d), but the effect was not statistically significant in the final model. No multicollinearity was detected for any of the covariates in the adjusted models (VIF < 5).

**Table 3.** Associations of the (CADASIL diagnosis x standardized ePVS volume) interaction term with imaging and cognitive measures. Baseline and adjusted models are presented. Abbreviations: BPF = Brain Parenchymal Fraction, CADASIL = Cerebral Autosomal Dominant Arteriopathy with Subcortical Infarcts and Leucoencephalopathy, CDRBox = Clinical Dementia Rating Sum-of-Boxes, MMSE = Mini Mental State Examination, Rey10m = Rey-Osterrieth Complex Figure Test delayed recall at 10 minutes, TRAILB = Trail B Test completion time, WMH = White Matter Hyperintensity. *p-value < 0.05.

| Outcomes | Baseline models |  | Adjusted models for age, gender, education |  |
| --- | --- | --- | --- | --- |
| | $\beta$ (95% CI) | p value | $\beta$ (95% CI) | p value |
| WMH volume (scaled) | 0.57<br>(0.12 to 1.02) | **0.02\*** | 0.45<br>(0.03 to 0.87) | **0.03\*** |
| BPF | -0.03<br>(-0.06 to 0.002) | 0.07 | -0.02<br>(-0.05 to 0.0007) | 0.06 |
| CDRBox | 0.52<br>(0.08 to 0.95) | **0.02\*** | 0.39<br>(0.01 to 0.77) | **0.04\*** |
| MMSE | -1.49<br>(-2.81 to -0.16) | **0.03\*** | -1.06<br>(-2.43 to 0.31) | 0.13 |
| TRAILB | 43.23<br>(-43.52 to 129.97) | 0.32 | 9.60<br>(-74.41 to 96.62) | 0.82 |
| Rey10m | -0.10<br>(-1.65 to 1.45) | 0.90 | 0.06<br>(-1.60 to 1.72) | 0.94 |

**Figure 1.**
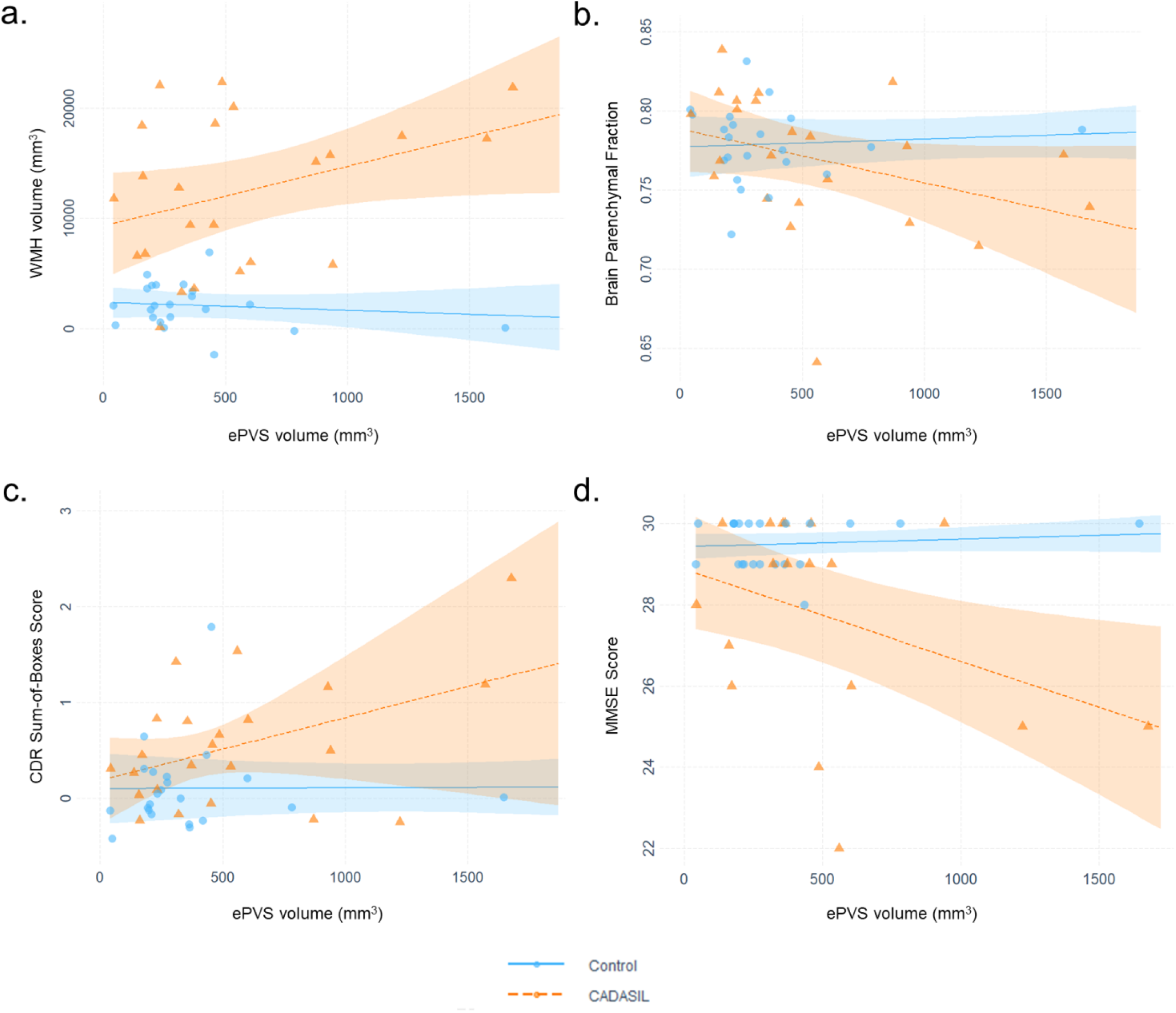
Interaction plots with lines fitted for the CADASIL diagnosis and ePVS interaction models. Points represent partial residuals and shaded areas 95% confidence intervals. Graphs illustrate the results of a. adjusted model with WMH volume as outcome b. adjusted model with BPF as outcome c. adjusted model with CDR Sum-of-Boxes score as outcome d. baseline model with MMSE score as outcome. The point of one control participant with relatively large ePVS volume is not shown due to size constraints. BPF = Brain Parenchymal Fraction, CDR = Clinical Dementia Rating, ePVS = Enlarged Perivascular Space, MMSE = Mini Mental State Examination, WMH = White Matter Hyperintensity

### Association between proteins from plasma proteomics and ePVS burden

We hypothesized that these associations may be capturing interesting molecular mechanisms of disease in CADASIL. We proceeded to testing associations of plasma protein levels, obtained through the SomaScan 7k assay, with the clinical measures of interest in the CADASIL group. 194 SomaScan probes, which corresponded to 190 unique proteins, were significantly associated with ePVS volumes, out of which 145 were in positive and 45 were in the negative direction. We then assessed protein-protein interaction networks (PPI) using the STRING database. For the positively associated proteins, 95/145 were functionally connected, and this network had a higher number of interactions than expected by chance (number of edges: 198; expected number of edges: 118; PPI enrichment p-value: 1.46e-11) (Fig. 2a). GO analysis of the biological processes revealed that the majority of enriched terms after FDR adjustment involved immune system processes, as well as neuronal axon function (Fig. 2b). We undertook a similar approach for negatively associated proteins, with 11/45 being connected significantly (number of edges: 8; expected number of edges: 4; PPI enrichment p-value: 0.03) (Fig. 2c). The biological process GO terms that were enriched for this subset mostly concerned lipid metabolism. Interestingly, the term for artery morphogenesis was also enriched (Fig. 2d).

**Figure 2.**
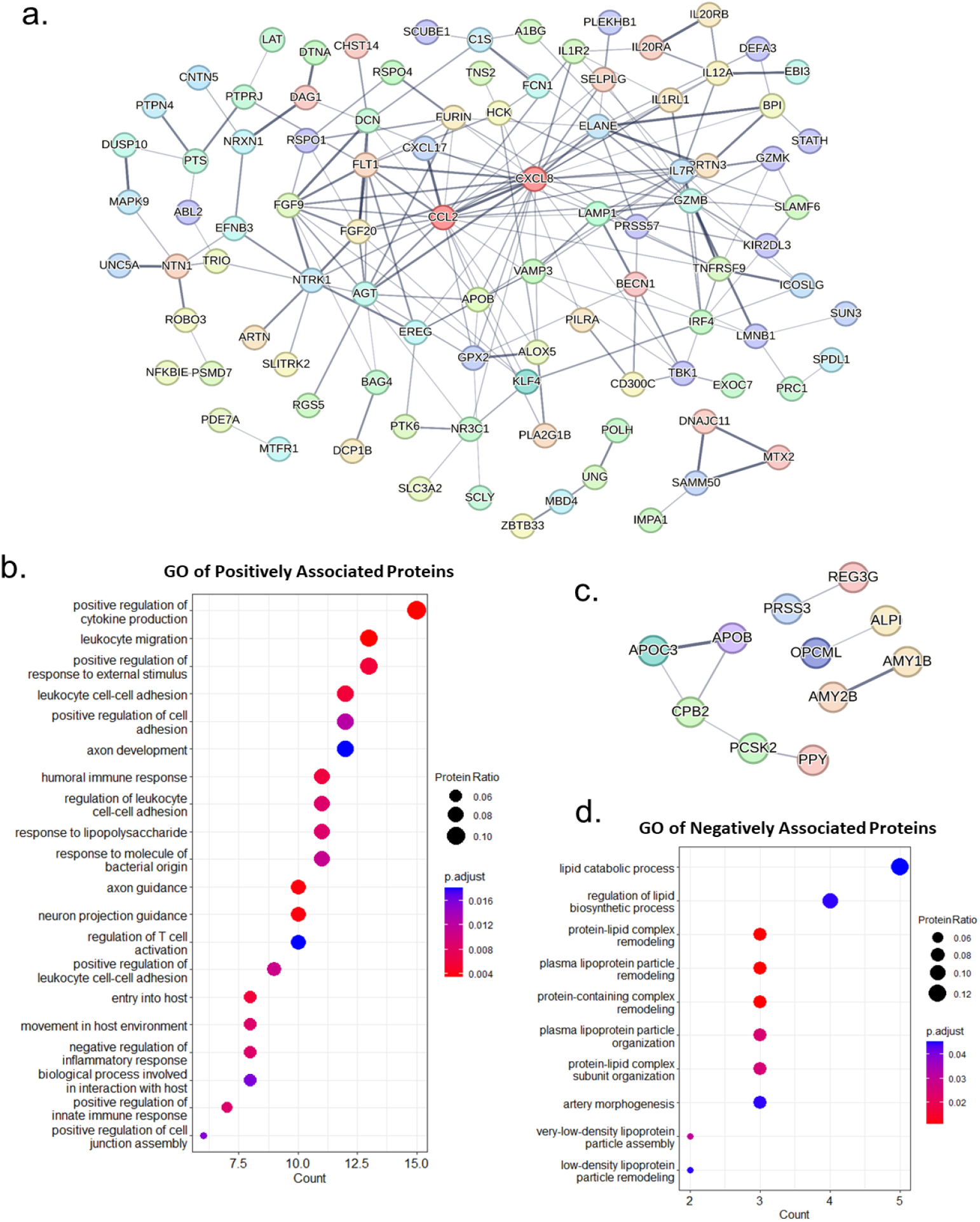
Protein-protein interactions and gene ontology analysis of proteins associated with ePVS volume. a. Interaction network between positively associated proteins. The proteins with most interactions (CCL2 - 24 edges, CXCL8 - 23 edges) are colored in dark red. b. Dot plot presenting the result of BP GO analysis for the all the positively associated proteins. c. Interaction network between positively associated proteins. d. Dot plot presenting the results of BP GO analysis for the all the negatively associated proteins. Intensity of edges in the interactions plots denote confidence. Proteins with no interactions are not shown. For the GO analyses, only significant terms (adjusted p-value < 0.05) were included. Protein Ratio is defined as the count of proteins in the term to the total number of proteins analyzed. Abbreviations: BP = biological process, GO = gene ontology.

### Associations between ePVS-associated proteins and clinical measures

We noticed that 2 proteins were highly connected in our PPI networks, thereby acting as protein network hubs: C-C motif chemokine ligand 2/ monocyte chemoattractant protein 1 (CCL2/MCP-1: 24 edges) and CXC motif chemokine ligand 8/interleukin-8 (CXCL8/IL-8: 23 edges) (Fig. 2a). We then examined possible associations of these hub proteins with our clinical measures of interest. Increased levels of CXCL8/IL-8 were significantly associated with increased ePVS (β = 2.44; 95% CI = 1.23 to 3.64; p = 0.0004) and WMH burden (β = 42.86; 95% CI = 3.80 to 81.92; p = 0.03) (Fig. 3a). An increase in CCL2/MCP-1 levels was also associated with increased ePVS levels (β = 2.77; 95% CI = 0.73 to 4.81; p = 0.01), Trail B completion time (β = 0.76; 95% CI = 0.45 to 1.07; p = 0.00007) and decreased BPF (β = -0.0007; 95% CI = -0.001 to -0.0003; p = 0.0003) and MMSE scores (β = -0.02; 95% CI = -0.04 to -0.007; p = 0.009) (Fig. 3b). Delayed Rey - Osterrieth Complex Figure test score was also marginally decreased (β = -0.02; 95% CI = -0.03 to 0.0006; p = 0.06).

**Figure 3.**
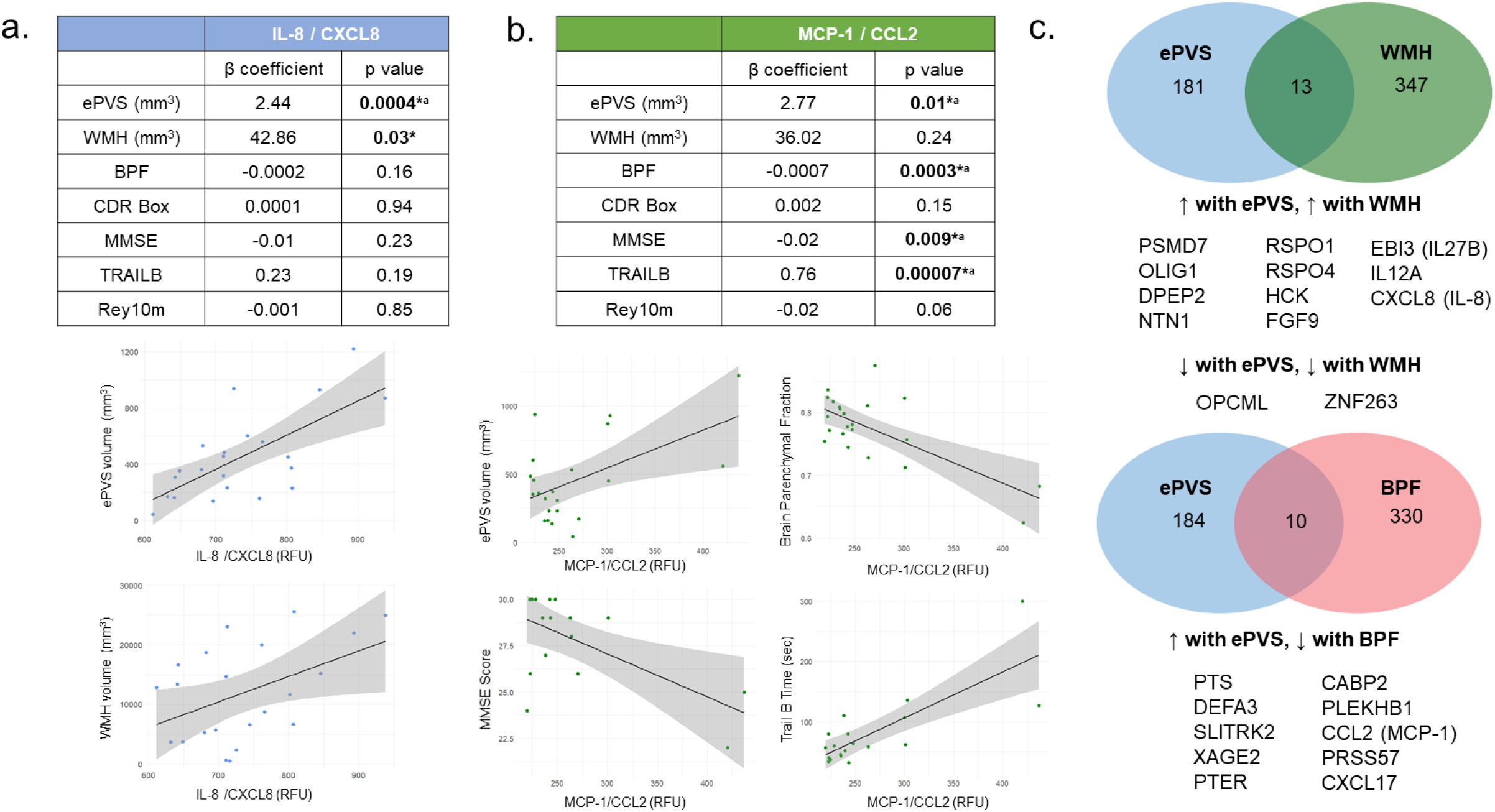
Protein expression is associated with characteristic findings in CADASIL patients. a. Linear models with IL-8/CXCL8 as predictor and the corresponding graphs for the models with ePVS and WMH volume as outcomes. b. Linear models with MCP-1/CCL2 as predictor and the corresponding graphs for the models with ePVS volume, BPF, MMSE score and Trail B completion time as outcomes. c. Overlap of proteins associated with specific neuropathological findings. Note that there was no protein associated with all three measures simultaneously. Abbreviations: BPF = Brain Parenchymal Fraction, CDRBox = Clinical Dementia Rating Sum-of-Boxes, MMSE = Mini Mental State Examination, Rey10m = Rey-Osterrieth Complex Figure Test delayed recall at 10 minutes, TRAILB = Trail B Test completion time, WMH = White Matter Hyperintensity. *p-value < 0.05, ^a^significant after Benjamini-Hochberg FDR correction.

WMH scores were not significantly associated with CCL2/MCP-1 expression (Fig. 3b). This result prompted us to test whether there was a shared molecular marker of all three relevant patho-anatomical measures, namely ePVS, WMH and BPF. We performed linear regression models with our proteomics data as predictors and WMH and BPF as outcomes. We found significant associations with 360 and 340 SomaScan probe levels respectively. For WMH, 13 proteins overlapped with the ePVS-associated proteins. The directionality of associations was the same for both ePVS and WMH volume (Fig. 3c). Similarly, 10 proteins were shared between the BPF- and ePVS-associated protein sets (Fig. 3c). Increased levels of all 10 proteins predicted increased ePVS volume and decreased BPF, as expected by the natural history of CADASIL. However, there was no common associated protein between all measures, suggesting degrees of biological separation between WMH volumes and brain atrophy in CADASIL. Alternatively, WMH provides a history of the brain, with signal change remaining on MRI chronically. Therefore, it is not as dynamic a measure of disease progression.

## Discussion

To date, few studies have investigated the association of MRI-visible ePVS burden with clinically meaningful outcomes in CADASIL. In this work we used a semi-automated approach to quantify ePVS load in a deeply phenotyped CADASIL cohort and rigorously tested associations between ePVS volume and clinically relevant outcomes. Despite there being no significant difference in ePVS volumes between CADASIL and control participants, ePVS volume was significantly associated with increased WMH volumes, brain atrophy, and declined brain function only in the CADASIL group. Additionally, we investigated plasma proteomic associations of ePVS, which were predominantly involved in inflammatory processes and lipid metabolism.

While ePVS have been shown to increase with aging^25^, the relationship of ePVS with clinically significant measures of brain disease remains inconclusive. Volume of ePVS *per se* may not always be an indicator of underlying disease. Higher load of ePVS can be found in healthy adolescents, mostly in the frontal and parieral white matter^26^. This phenomenon could explain the lack of a difference in mean ePVS volumes between our groups. However, in the case of cSVD, ePVS volumes have been shown to be increased in patients with vascular-related cognitive impairment in comparison to other types of dementia and healthy individuals^27^, and in patients with lacunar versus large vessel strokes^28^.

In CADASIL, four studies have examined associations between clinical measures and ePVS volumes. All studies are limited by their use of visual counting methods and Likert-based categories of MRI-visible ePVS burden. The first study compared patients with few ePVS to patients with increased ePVS counts, and found no differences in MMSE or modified Rankin score (mRs) ^29^. In a larger study that followed a similar approach, the group of individuals with CADASIL demonstrated higher burden of ePVS and worse performance on Mattis Dementia Rating Scale (MDRS), MMSE, and lower mRS than individuals with low burden of ePVS, after adjusting for age, gender, and all relevant imaging markers of pathophysiology^30^. Interestingly, in the high ePVS burden group, temporal and subinsular ePVS volumes were associated with increased WMH volume, and increased white matter ePVS was marginally associated with decreased BPF ^30^. Another study found an association of ePVS in the centrum semiovale with backwards digit span (BDS), a measure of executive function, but no associations with other measures of cognition ^31^. Lastly, another group found that regional ePVS burdens in the anterior temporal lobe were associated with increased BFP, but regional ePVS were not associated with mRS score^32^. Likert-based categorization of patients can confound results due to the variability in individual ePVS volumes. Future application of automated methods for ePVS quantification, similar to the one we implemented, can increase the reliability and comparability between studies and contribute to the standardization of measurement.

We show that WMH is associated with ePVS volume in CADASIL. WMH volume has been shown to be associated with ePVS in patients with cSVD and healthy individuals^25, 28^. Additionally, basal ganglia ePVS in healthy individuals have been associated with measures of increased BBB permeability^33^. A post-mortem study in brains from CADASIL patients identified significant fibrinogen extravasation in WMH forming around ePVS, but not in in WMH without other pathology, suggesting that ePVS burden in CADASIL and the surrounding WMH may indicate BBB leakage^34^. Moreover, from mechanistic studies, fibrinogen has been identified as a trigger of neuroinflammation and synaptic degeneration^35^. This is reflected in the associations we found with plasma proteomic markers of immune activation, inflammation and axonal remodelling.

We demonstrated an association between ePVS volumes and brain atrophy, represented by decreased BPF, in CADASIL. A similar finding has been noted in various brain disorders, such as lacunar strokes^36^ and multiple sclerosis^37^. A neuropathological study in 4 CADASIL patients provided experimental evidence that markedly increased ePVS burden in the basal ganglia, termed *status cribosum*, was connected with cortical neuronal apoptosis and axonal degeneration^38^.

Regarding cognitive measures, our study reinforces previous findings that increased ePVS volumes may be associated with decline in measures of functionality (CDR Box Score) and general cognition (MMSE Score)^30^. A study on cSVD showed that decreased diffusion tensor image analysis along the perivascular space (DTI-ALPS) scores were associated with decreased MMSE scores, and DTI-ALPS mediated the relationship between WMH volume and episodic memory^39^. The inclusion of education in our models dampened the relationship with MMSE, possibly reflecting cognitive reserve, or inadequate power in models. The other cognitive measures studied of processing speed, executive function, and visual memory were chosen based on prior observations that processing speed and executive function deteriorate early in CADASIL^40, 41^, whereas delayed memory and visuospatial abilities may exhibit decline at late stages of the disease, but not consistently across patients^40, 41^. Furthermore, processing speed decline in patients with CADASIL has been associated with brain atrophy^7^. In our study, processing speed did not associate with ePVS volumes in the CADASIL group, but we noted an association in the combined group. This is concordant with prior studies of sporadic cSVD, reporting on association of ePVS volumes with decreased processing speed and executive function scores^42^.

To connect the clinical associations of ePVS with disease pathophysiology, we investigated possible associations of ePVS volumes with plasma protein levels. Immune activation pathways, specifically leukocyte adhesion and migration, were significantly enriched in our analyses. Circulating markers of inflammation (IL-6, fibrinogen and C-reactive peptide)^43^, neutrophil counts and increased neutrophil-to-lymphocyte ratio^44^ have all been associated with increased ePVS burden in the general population. We identified 2 chemokines central to the interaction networks of ePVS-relevant proteins, CXCL8/IL-8 and CCL2/MCP-1. CXCL8/IL-8, produced, among others, by microglia and endothelial cells^45^, and CCL2/MCP-1, are both neutrophil and monocyte chemo-attractants^45, 46^. In individuals with normal cognition or mild cognitive impairment, these cytokines have been implicated in cognitive decline^45^ and age-related memory loss ^46^. CXCL8/IL-8 has further been associated with WMH burden ^45^ and increased ischemic stroke risk^47^. A mechanism that has been suggested to mediate CCL2/MCP-1 effects on brain injury is activation of microglia after acute damage and CD8^+^ T-cell recruitment^48^, which was enriched in our GO-term analysis. Interestingly, a recent article described CCL2/MCP-1 as being up-regulated by aberrant Notch signaling in patients with nonalcoholic steatohepatitis, driving liver monocyte infiltration and fibrosis^49^. The above suggest that the observed phenotypic associations in CADASIL could be of immunovascular origin, however targeted mechanistic studies are needed to prove this.

Our study’s strengths include the age and gender-matching, the deep phenotyping of individuals and the multi-modal models. To our knowledge, this is the first study to connect clinical correlates of ePVS in CADASIL with molecular markers in peripheral blood. With regards to limitations, this study may have been underpowered to detect some associations. We partially circumvented this issue by the use of continuous models. In addition, static or cross-sectional investigations may not adequately capture ePVS abnormalities, which have been shown to be dynamic, with diurnal change^50^. Also, we did not account for specific mutations. Patients with CADASIL mutations in EGFr 1-6 have been shown to harbor greater ePVS volume in the ATL and subinsular regions than patients with in EGFr 7-34 mutations^32^. Furthermore, future studies can investigate whether there is value in quantifying regional ePVS volumes in CADASIL. In the general population, regional ePVS load has been associated with specific risk factors ^25^.

We found that greater ePVS volumes were associated with increased imaging measures of brain injury and degeneration, worse cognitive outcomes, and markers of immune cell recruitment and inflammation in CADASIL. Whether they are representative on a microscopic level of inflammatory processes at the BBB, poor glymphatic flow or other etiology remains to be confirmed. Additional studies to further validate our reported findings and establish ePVS as a relevant marker of disease progression in CADASIL would be of great value.

## Glossary

BBB: blood-brain barrier
BPF: brain parenchymal fraction
CADASIL: Cerebral Autosomal Dominant Arteriopathy with Subcortical Infarcts and Leucoencephalopathy
CCL2/MCP-1: C-C motif chemokine ligand 2/ monocyte chemoattractant protein 1
CDRBox: Clinical Dementia Rating sum-of-boxes
CMBs: cerebral microbleeds
CSF: cerebrospinal fliud
cSVD: cerebral Small Vessel Disease
CXCL8/IL-8: CXC motif chemokine ligand 8/interleukin-8
EGFr: epidermal growth factor–like repeats
ePVS: enlarged perivascular spaces
FLAIR: Fluid Attenuated Inversion Recovery
MMSE: Mini Mental State Examination
Rey10m: Rey-Osterrieth Complex Figure Test delayed recall at 10 minutes
TIV: total intracranial volume
TRAILB: Trail B Test completion time
VCID: vascular cognitive impairment and dementia
WMH: white matter hyperintensity.

## Acknowledgements

The authors thank all patients and families for their generous participation in research, without whom this work would not have been possible. The authors also thank staff and investigators at the UCSF Memory and Aging Center that make excellent clinical research possible.

